# A field test for frequency-dependent selection on mimetic colour patterns in *Heliconius* butterflies

**DOI:** 10.1101/005249

**Authors:** Patricio A. Salazar, Martin Stevens, Robert T. Jones, Imogen Ogilvie, Chris D. Jiggins

**Author notes:** Corresponding author: Patricio A. Salazar Current Address: Centro para la Investigaciόn de la Biodiversidad y el Cambio, Climático - BioCamb, Universidad Tecnolόgica Indoamérica, Machala y Sabanilla, 1703175 - Quito, Ecuador /.

## Abstract

Müllerian mimicry, the similarity among unpalatable species, is thought to evolve by frequency-dependent selection. Accordingly, phenotypes that become established in an area are positively selected because predators have learnt to avoid these forms, while introduced phenotypes are eliminated because predators have not yet learnt to associate these other forms with unprofitability. We tested this prediction in two areas where different colour morphs of the mimetic species *Heliconius erato* and *H. melpomene* have become established, as well as in the hybrid zone between these morphs. In each area we tested for selection on three colour patterns: the two parental and the most common hybrid. We recorded bird predation on butterfly models with paper wings, matching the appearance of each morph to bird vision, and plasticine bodies. We did not detect differences in survival between colour morphs, but all morphs were more highly attacked in the hybrid zone. This finding is consistent with recent evidence from controlled experiments with captive birds, which suggest that the effectiveness of warning signals decreases when a large signal diversity is available to predators. This is likely to occur in the hybrid zone where over twenty hybrid phenotypes coexist.

## INTRODUCTION

Müllerian mimicry evolves because two or more unpalatable species gain mutual advantage from advertising a common display. In particular, mimicry reduces the number of individuals that need to be killed per species before a predator learns to identify them as distasteful (Joron & Mallet, 1998; Mallet & Joron, 1999; Sherratt, 2008). The species pair *Heliconius erato* Linnaeus, 1758 and *H. melpomene* Linnaeus, 1758 is a famous case of Müllerian mimicry. These distantly related species are widely distributed in Central and South America, and despite their great geographical variation in wing colour pattern they show almost identical colour patterns wherever they co-occur. The persistence of numerous mimetic races of *H. erato* and *H. melpomene* seems paradoxical, because these intra-specific variants can easily hybridize in nature, a process that promotes the mixing of these divergent populations into a single hybrid pool. Furthermore, frequency-dependent selection for mimicry should favour the existence of one single colour pattern. Therefore, in order to maintain their colour pattern distinctiveness, phenotypes that have reached a high frequency in a particular area are expected to be positively selected, while novel colour morphs should tend to be eliminated. Frequency-dependent selection in *Heliconius* is thought to be mainly caused by bird predators, as birds have excellent colour vision (Cuthill, 2006) and are well known to discriminate tasteful from distasteful prey (e.g. Brower, *et al*., 1971; Langham, 2004; Ihalainen *et al*., 2007).

Experiments with captive birds have shown that known *Heliconius* predators avoid familiar colour patterns, but repeatedly attack novel forms (Langham, 2004), and field studies using mark recapture experiments have revealed that mimetic (locally established) colour patterns have higher survival than non-mimetic (introduced) patterns (Benson, 1972; Mallet & Barton, 1989; Kapan, 2001). However, there is no direct evidence that these survival differences are caused by birds (Mallet & Barton, 1989). Moreover, mark-recapture studies do not tease apart the potentially confounding effects of local adaptations, other than colour pattern, on the survival of trans-located individuals (Mallet & Barton, 1989; Kapan, 2001). In this study, we attempted to overcome these limitations by using artificial prey, a method that had successfully been used to test for aposematism or mimicry in various organisms (e.g. Brodie III, 1993; Comeault & Noonan, 2011; Valkonen *et al*., 2011), including *Heliconius* and other Lepidoptera models (Cuthill *et al*., 2005; Stevens *et al*., 2008; Finkbeiner *et al*., 2012; Merrill *et al*., 2012). Our *Heliconius* models were created with paper wings that resembled real butterflies to bird vision, and plasticine bodies that allowed us to identify predation events by the marks left on the plasticine (Figure 1). Moreover, because the only difference between models is their colour pattern, this method allowed us to test for colour pattern effects while controlling for the effects of potentially confounding factors that exist when studying living butterflies (e.g. toxin levels, behavioural differences, or physiological adaptations to different thermal environments).

**Figure 1.**
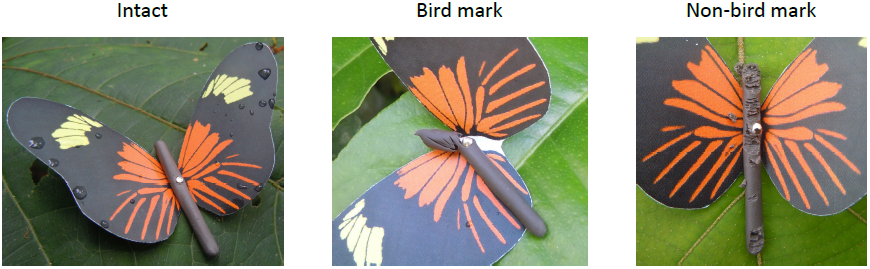
Photographs of intact and marked models. Bird marks were easily identified because they typically consisted of one single—usually triangular—dent, whereas non-bird marks consisted of many scratches.

In addition, we investigated the pattern of bird attacks on the butterfly models by recording the location of each attack on the plasticine body. We were interested in this aspect of bird predatory behaviour because there is evidence that birds tend to target their prey’s head. For instance, several butterfly species, especially Lycaenids, have evolved head mimicking structures in the posterior end of their hind wings, which have been shown to deflect bird attacks away from the head (Wourms & Wasserman, 1985). In our case, confirming that bird marks tend to occur on the butterflies’ head would further support the expectation that the artificial (and real) prey are preyed upon by birds.

In this study, we focused on the *H. erato* and *H. melpomene* mimicry system in eastern Ecuador, South America. Here, in the upper Pastaza river valley, above 1100 m in elevation, the endemic colour pattern race *H. e. notabilis* coexists with its co-mimic *H. m. plesseni*. Further east into the Amazonian lowlands, below 500 m, a different mimetic pair of *H. erato* and *H. melpomene* races occurs (*H. e. lativitta* and *H. m. malleti*). At mid-elevation, between 500 and 1100 m, the races within each species hybridize across an approximately 38km-wide zone (Figure 2A). Models of three *H. erato* colour patterns were exposed to predation in three zones: the two inhabited by pure races and the hybrid zone (Figure 2B). In each zone we exposed models of the locally established or most frequent form, as well as two introduced colour patterns (one of a pure race and one hybrid; Table 1). Contrary to predictions, no differences in attack rates were detected between the different aposematic patterns tested. However, higher overall predation rates were recorded in the hybrid zone, which is characterised by a diverse prey community, and a strong tendency for avian predators to target the prey’s head was observed across all studied zones.

**Figure 2.**
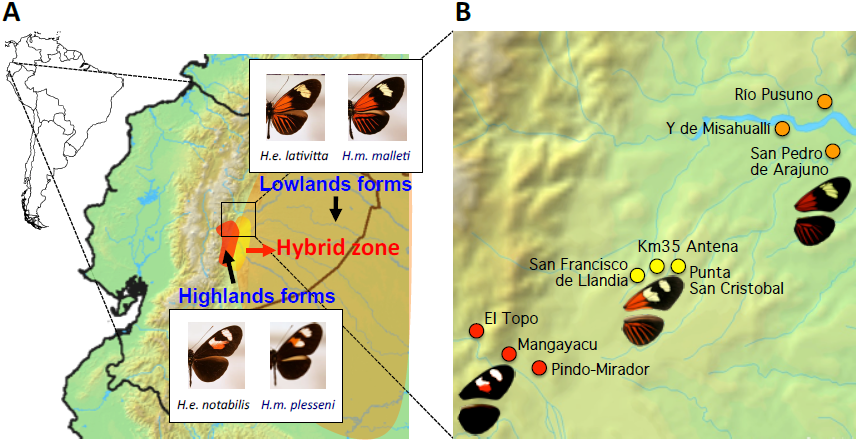
(A) Geographic location of the study area, and mimicry between *Heliconius erato* and *H. melpomene*. Hybridization occurs between races within each species: *H. e. lativitta* hybridizes with *H. e. notabilis*, and *H. m. malleti* hybridizes with *H. m. plesseni*. (B) Study sites (red, orange, and yellow circles) and locally established or most abundant colour patterns of *H. erato*.

**Table 1.**
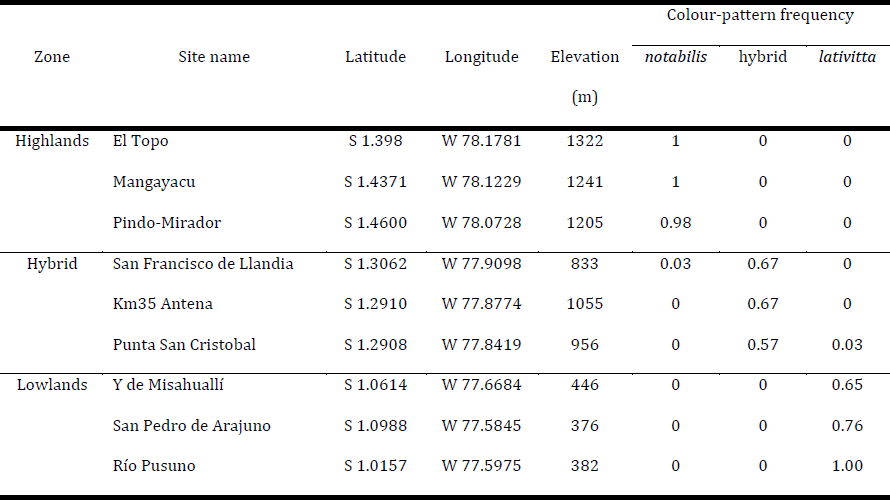
Location of experimental sites in Eastern Ecuador and frequency of each tested colour-pattern per experimental site (colour-pattern frequencies do not add up to 1 within sites, because other hybrid colour-patterns—less abundant than that tested in this study—could have been present at any particular site; see Salazar 2012, Chapter 3, for details).

## MATERIALS AND METHODS

### Prey stimuli

Artificial prey were paper ‘butterflies’ printed with specific patterns to resemble the different pure and hybrid forms tested in this study, along with a plasticine body (Figure 1). Stimuli wings were made from waterproof paper (HP LaserJet Tough Paper, Palo Alto, USA) printed with the different colour patterns of the real butterflies using a Hewlett Packard LaserJet 2605dn printer at 300 dpi. The colour patterns used were based on digital photographs of real butterfly specimens (four specimens per colour-pattern), taken with a Fujifilm IS Pro UV-sensitive digital camera with a quartz CoastalOpt ultraviolet (UV) lens (Coastal Optical systems). It is well known that birds see a wider colour spectrum than humans do (Cuthill, 2006), and stimuli printed from uncalibrated printers rarely correspond closely to real object colours, especially to non-human vision. Colours printed on the paper wings were therefore calibrated so that they stimulated the photoreceptors of potential avian predators in a similar way to real wings. Specifically, appropriate colours were selected so that when printed they produced photon catch values for each cone type that fell within the range of values for the corresponding colour patches on the real butterflies (as with past work on camouflage; e.g. Cuthill *et al*., 2005; see also Supplementary Material). All colours present on the studied phenotypes were closely reproduced, except for the white patches of the highlands’ race (*H. e. notabilis*), whose reflectance in the UV range it was not possible to reproduce accurately (Figure 3; Supplementary Material, Figure S1).

**Figure 3.**
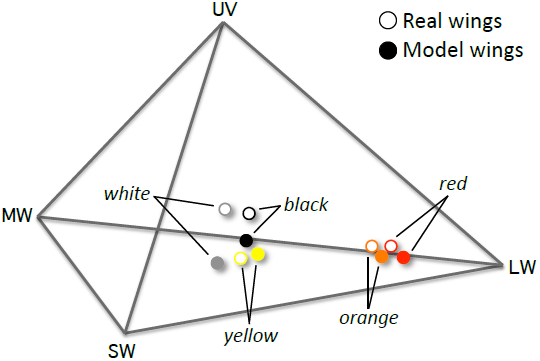
Location of colour patches of real wings (open circles) and paper models (filled circles) in the tetrahedral colour space defined by the photon-catch values for each cone type of the bird retina: ultraviolet (UV), short (SW), medium (MW) and long (LW) wavelength sensitive cones. Notice that all colours in the models were reproduced very close to those of real wings. The only exception was the white patches of the *H. erato* highlands’ race (*H. e. notabilis*), whose reflectance in the UV range it was not possible to reproduce (see Supplementary Material for details).

### Experimental design for frequency-dependent selection test

We conducted field predation experiments with the artificial butterflies using a modified version of a protocol well established for testing the survival value of camouflage and eyespot markings with artificial prey (e.g. Cuthill *et al*., 2005; Stevens *et al*., 2008). Here, prey items are pinned out at low density along non-linear transects and monitored for predation. We applied a fully crossed factorial design, with two study factors (colour pattern and zone), each having three levels of variation (3 colour patterns x 3 zones). If the hypothesis of frequency-dependent selection was to be supported, each colour pattern was expected to survive best in the zone where it reaches the highest frequency, while introduced (non-mimetic) colour patterns should have lower survival. We carried out experimental trials in three replicate sites per zone (Figure 2B), each separated by at least 4 km. In each site, 80 models per pattern were set and exposed to predation for 72 hours. One model per pattern was placed every 10-13 m along accessible pathways in areas where the studied *Heliconius* species were observed. Models were checked for signs of attack every 24 hours. In order to optimize the methodology, we first carried our pilot experiments between December 2009 and March 2010, and the actual test for frequency-dependent selection took place between July and September 2010. Details of how potential confounding factors were controlled for are described in the Supplementary Material.

### Analysis of predation marks data

We tested the hypothesis of frequency-dependent selection as part of a set of competing hypotheses that could explain the predation marks data. We followed a model selection approach based on the Akaike information criterion (AIC; Burnham & Anderson, 2002). The data consisted of time-to-mark observations for every model that was placed in the field. Models that remained intact for the full 72 h of the experimental trials were considered right-censored (i.e. the time to mark is unknown, but is certainly longer than 72 h). Data for marked models, in contrast, were treated as interval-censored (e.g. when a model was found marked at 24 h, it is known that the mark occurred at some point in the interval 0-24 h). Treating time-to-mark data as interval-censored allowed us to use of all observations in the analysis, including those in which a model was not found in one or more censuses but was re-sighted later; in this latter case, the interval when the mark is known to occur is longer than the standard 24 h, but remains informative.

As first step in the analysis, we developed a set of competing hypotheses that could explain the information contained in the data, and formalized them as statistical models. The experimental design included the possible effect of the following factors: colour pattern, zone and experimental site (nested within zone). Since the hypothesis of interest predicts that each tested colour pattern should be the least attacked in the zones where they constitute the most frequent phenotypes, we expected the statistical interaction between colour pattern and zone to be strongly supported by the data. Thus, the final set of competing statistical models included different combinations of the experimental design factors, including the interaction between colour pattern and zone.

Statistical models were fitted in Minitab ® (Minitab Inc., 2010), using the ‘Regression with life data’ tool. In order to model the hazard function for the time-to-mark data (i.e. the function that describes the probability of a butterfly model being marked through the course of the experiment), we assessed the goodness-of-fit of several probability distributions before fitting any of the competing models. We decided to use different probability distributions for bird and non-bird marks. For bird marks, we assumed an exponential distribution—which presupposes a constant hazard rate—because, given the small number of bird marks recorded, there was not enough information in the data to justify a more complex function. For non-bird marks, in contrast, we assumed a Weibull distribution—which can fit either increasing, constant, or decreasing hazard rates—because the number of non-bird marks recorded was large enough to give signs of variation in hazard rates for butterfly models affected by different factors, and the fitting algorithm for models that assumed the more restrictive exponential distribution rarely converged.

Once the statistical models were fitted, we computed the AIC for each of the competing models, and ranked the models accordingly. The relative support in the data for each competing model was assessed on the basis of their Akaike weights (*w*_*i*_), and the evidence ratios obtained by dividing the weight of the best-ranked model by the weight of each competing model (*w*_1_/*w*_*i*_). Akaike weights range from zero to one, and can be interpreted as an approximate probability of each model being the actual best-ranked model, if the model selection exercise would be repeated with a different but equivalent data set (Burnham & Anderson, 2002). Finally, in order to visualize the survival patterns of the butterfly models through the course of the experimental trials, we reconstructed survival curves based on non-parametric Turnbull estimates of survival probabilities for interval-censored data (Turnbull, 1976; Minitab Inc., 2010).

### Analyses of marked body sections

In order to test the hypothesis that birds were more likely to attack the head of the plasticine bodies rather than any section at random, we made all plasticine bodies the same length (3 cm) and recorded the body section (or sections) where attack marks occurred. We considered three equally sized sections: anterior, middle and posterior, and tested whether the frequency of marks occurring in one or more sections was larger than expected if all sections would be equally targeted, using a *G* test.

Several organisms marked the models. However, bird marks were easily distinguished from non-bird marks as they typically consisted of one single—usually triangular—dent, as opposed to many scratches (Figure 1). Moreover, we directly observed attacks from ants, grasshoppers and sweat bees in the field, and these explained the appearance of the majority of non-bird marks. We performed analyses for both bird and non-bird marks, as it is reasonable to expect different attack patterns for birds, which are visual predators, than for other organisms (e.g. insects) that most probably attacked the models without regard to the colour pattern they exhibited. Models marked by either birds or non-birds were taken away and not replaced.

## RESULTS

### Test for frequency-­dependent selection on colour patterns

We recorded 56 bird and 667 non-bird marks in a total of 2160 models exposed to predation. The low number of avian attacks compared to models probably reflects the fact that many birds will have learnt to avoid the real butterflies. Bird marks did not support the hypothesis of frequency-dependent selection for mimicry. None of the tested colour patterns survived best in the zones where they are the most abundant phenotypes (Figure 4A). Moreover, none of the statistical models that incorporate the interaction between colour pattern and zone were well supported. In fact, the simplest model containing the interaction parameter was ranked third, with an Akaike weight of 0.006, which is respectively eight and five times less supported than the first and second ranked models (Table 2). Furthermore, colour pattern is in general a poor predictor of the survival of butterfly models, as the survival curves for different colour patterns tend to overlap in all zones (Figure 4A), and the statistical models containing the factor colour pattern are not consistently ranked among the best supported (Table 2). The zone factor, in contrast, has a much stronger effect. The survival of all butterfly models, irrespective of colour pattern, is several times lower in the hybrid zone as compared to the parental zones (Figure 4B), and the first two ranked statistical models, which incorporate the zone factor, have a combined Akaike weight (*w*1 + 2) of 0.93. This implies approximately thirteen times more support for these first two models than for all other contested models (Table 2). Importantly, even though the first ranked model incorporates the parameter ‘colour pattern’ in addition to ‘zone’, the Akaike weights for each of the first two models are not very different. Thus, there is no evidence that one of the two best-ranked models has more support in the data than the other, and consequently the zone factor alone is a good predictor of the survival data.

**Figure 4.**
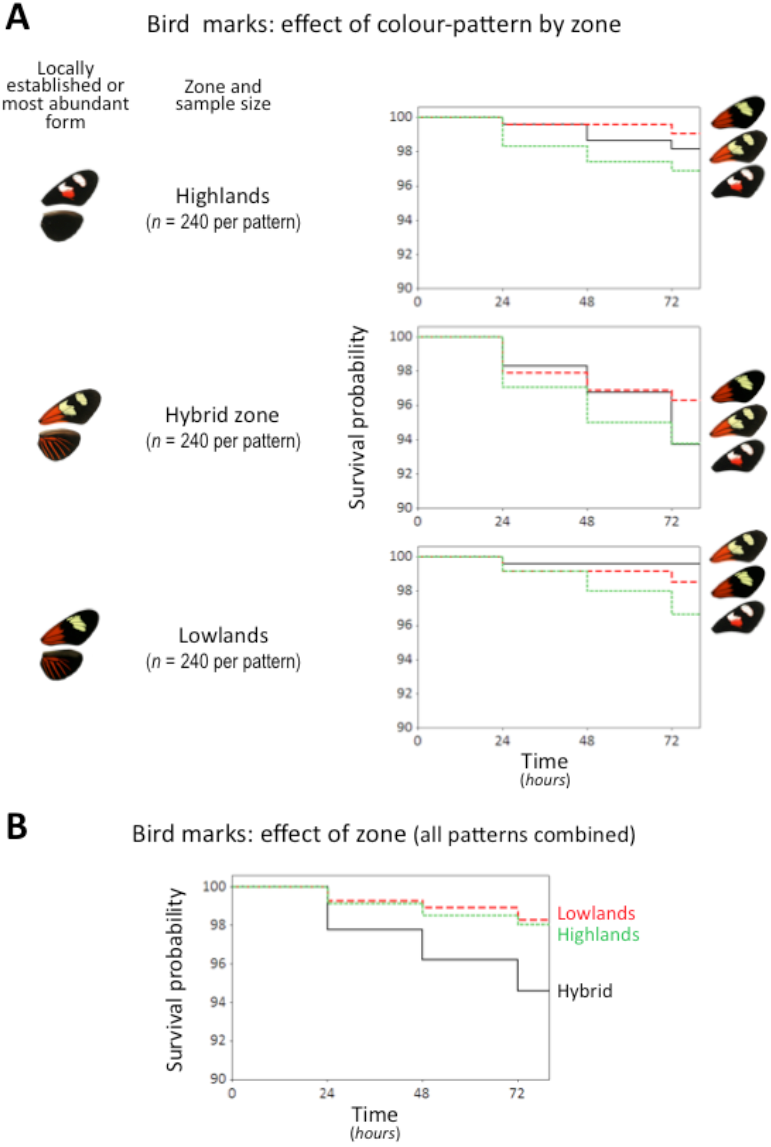
Non-parametric survival curves over time for bird marks: (A) effect of colour pattern in the survival of butterfly models within each zone, (B) effect of zone, disregarding colour pattern.

**Table 2.**
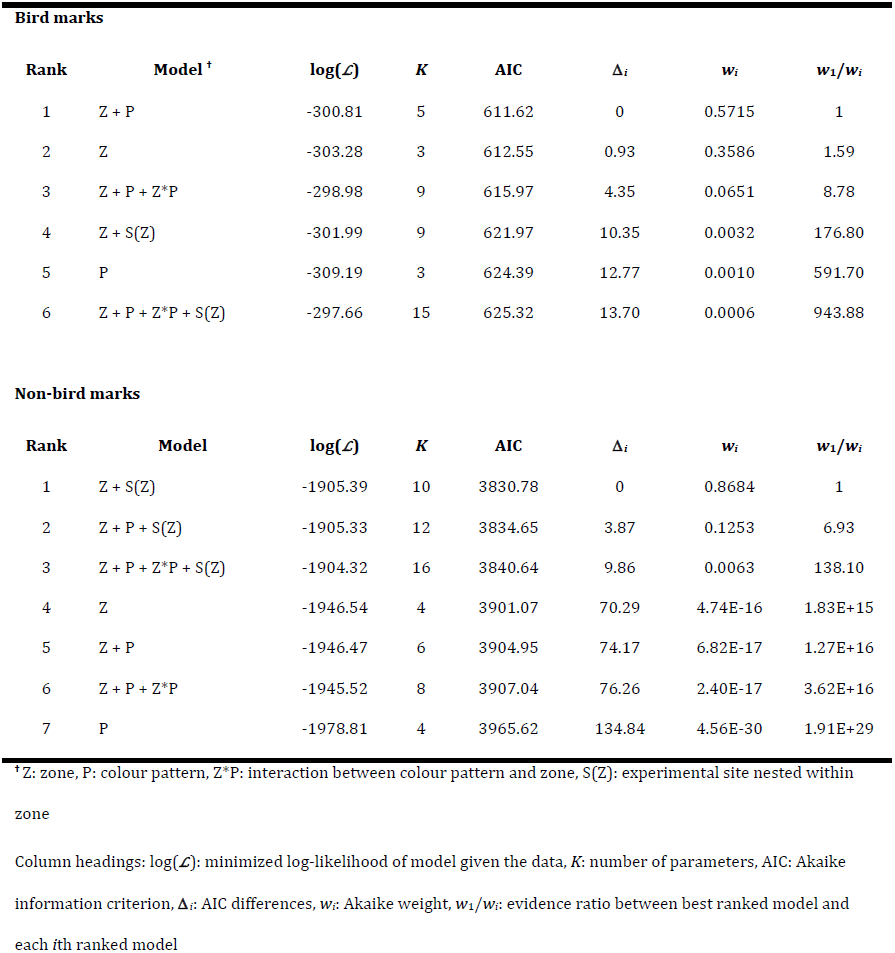
Statistical model comparison based on AIC.

| Bird marks |  |  |  |  |  |  |  |
| --- | --- | --- | --- | --- | --- | --- | --- |
| Rank | Model <sup>†</sup> | log($\mathcal{L}$) | $K$ | AIC | $\Delta_i$ | $w_i$ | $w_1/w_i$ |
| 1 | Z + P | -300.81 | 5 | 611.62 | 0 | 0.5715 | 1 |
| 2 | Z | -303.28 | 3 | 612.55 | 0.93 | 0.3586 | 1.59 |
| 3 | Z + P + Z*P | -298.98 | 9 | 615.97 | 4.35 | 0.0651 | 8.78 |
| 4 | Z + S(Z) | -301.99 | 9 | 621.97 | 10.35 | 0.0032 | 176.80 |
| 5 | P | -309.19 | 3 | 624.39 | 12.77 | 0.0010 | 591.70 |
| 6 | Z + P + Z*P + S(Z) | -297.66 | 15 | 625.32 | 13.70 | 0.0006 | 943.88 |
| Non-bird marks |  |  |  |  |  |  |  |
| Rank | Model | log($\mathcal{L}$) | $K$ | AIC | $\Delta_i$ | $w_i$ | $w_1/w_i$ |
| 1 | Z + S(Z) | -1905.39 | 10 | 3830.78 | 0 | 0.8684 | 1 |
| 2 | Z + P + S(Z) | -1905.33 | 12 | 3834.65 | 3.87 | 0.1253 | 6.93 |
| 3 | Z + P + Z*P + S(Z) | -1904.32 | 16 | 3840.64 | 9.86 | 0.0063 | 138.10 |
| 4 | Z | -1946.54 | 4 | 3901.07 | 70.29 | 4.74E-16 | 1.83E+15 |
| 5 | Z + P | -1946.47 | 6 | 3904.95 | 74.17 | 6.82E-17 | 1.27E+16 |
| 6 | Z + P + Z*P | -1945.52 | 8 | 3907.04 | 76.26 | 2.40E-17 | 3.62E+16 |
| 7 | P | -1978.81 | 4 | 3965.62 | 134.84 | 4.56E-30 | 1.91E+29 |

<sup>†</sup> Z: zone, P: colour pattern, Z\*P: interaction between colour pattern and zone, S(Z): experimental site nested within zone
Column headings: log( $\mathcal{L}$ ): minimized log-likelihood of model given the data, $K$ : number of parameters, AIC: Akaike information criterion, $\Delta_i$ : AIC differences, $w_i$ : Akaike weight, $w_1/w_i$ : evidence ratio between best ranked model and each $i$ th ranked model

For non-bird marks the results are different. Neither colour pattern nor zone (nor their interaction) are good predictors of the survival of butterfly models. All colour patterns showed similar survival within each zone (Figure 5A). However, attack rates were highly variable between sites within zones (Figure 5B). In fact, the parameter ‘experimental site’ (nested within zone) is the only well supported predictor of the non-bird marks data set. The first ranked model, which is the most reduced model containing the site parameter, has a much larger Akaike weight compared to all other candidate models (*w*_1_ = 0.87; Table 2).

**Figure 5.**
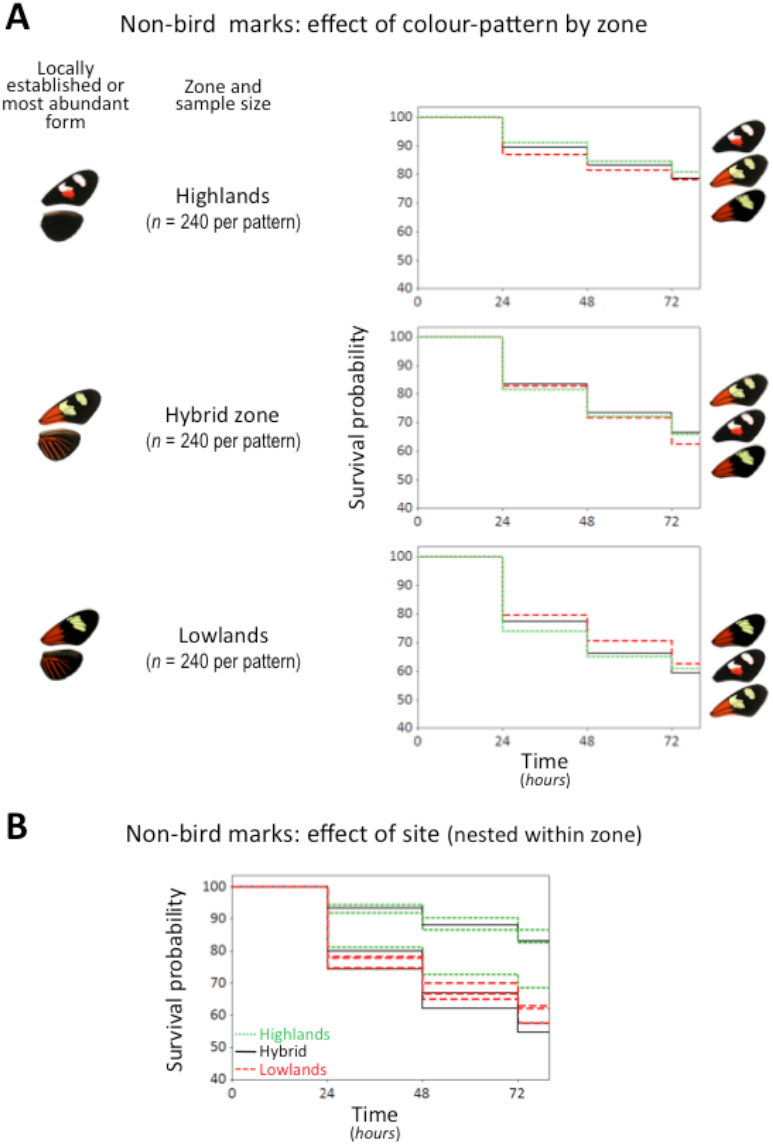
Survival curves for non-bird marks: (A) effect of colour pattern in the survival of butterfly models within each zone, (B) effect of experimental sites within zones (disregarding colour pattern). High variation in survival between sites blurs any potential difference in survival between zones.

### Analysis of marked body sections

There was a strong non-random distribution of bird attack marks on the body segments (*G*_6_ = 33.04, *p*< 0.001). Bird marks were more likely to affect the most anterior or the anterior + middle sections of the body, rather than any section or combination of sections at random. In contrast, non-bird marks were more evenly distributed across body sections (*G*_6_ = 562.89, *p*< 0.001, Figure 6).

**Figure 6.**
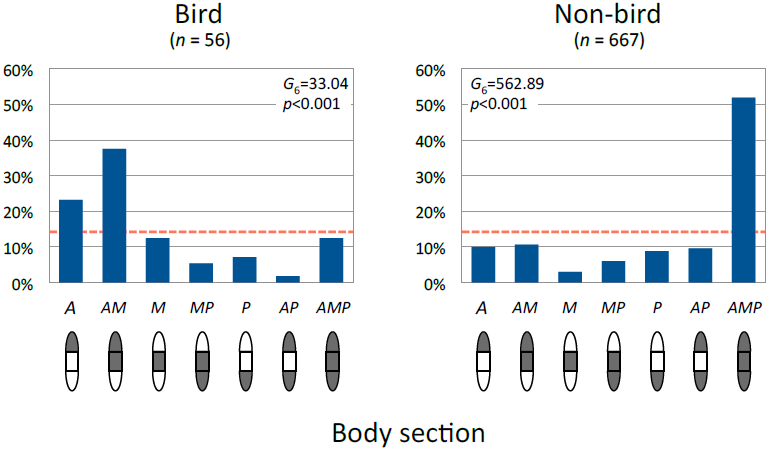
Frequency of bird and non-bird marks affecting a particular body section, or combination of sections. The body sections considered were A: anterior, M: middle and P: posterior. The red dashed line indicates the expected frequency if each section or combination of sections had been equally affected at random. Bird marks were more likely to affect either the anterior or anterior + middle sections. Non-bird marks, in contrast, were more likely to affect all body sections at the same time.

## DISCUSSION

### Is there frequency-­dependent selection for mimicry?

Unexpectedly, we found no evidence that bird attacks were associated with wing colour pattern. Furthermore, none of the locally established or most abundant colour patterns were favoured over introduced patterns (Figure 4A). Overall, and contrary to predictions, there was no statistical evidence that colour pattern had any effect on bird attack rates (Table 2). The inability to detect a colour pattern effect was probably due to a lack of statistical power, as the effect size of differential predation on colour patterns may be small, and therefore it would be necessary to gather a larger sample of informative data (i.e. bird marks) to have the power to detect such effects. Small colour pattern effects are perhaps not surprising, since all the tested colour patterns are aposematic and will therefore tend to be avoided. This is supported by the fact that we only recorded 56 avian attacks despite putting out a total of 2160 models. It is worth noticing, however, that this attack rate is within the range reported in other studies with artificial prey (Table 3), including at least one experiment with artificial *Heliconius* butterflies that did detect differences in bird predation associated with colour pattern (Merrill *et al*., 2012; Table 3). Hence, bird marks were not particularly infrequent in our study, but still not enough to detect any colour pattern effects that may exist. Importantly, the fact that we did not detect colour pattern effects does not mean that the hypothesis of frequency-dependent selection for mimicry is rejected. There is simply no evidence in our data that the locally established patterns were favoured in the zones were they are the most frequent phenotypes, but similarly there is no evidence that introduced colour patterns were favoured either.

**Table 3.** Summary of a sample of previous studies in which models were used to investigate natural selection on colour pattern.

| Publication | Organism | Hypothesis | Total<br>models | Bird marks |  | Non-bird marks |  | Exposure time<br>(h) | Colour<br>calibration | Support for<br>hypothesis |
| --- | --- | --- | --- | --- | --- | --- | --- | --- | --- | --- |
|  |  |  |  | No. | Freq. | No. | Freq. |  |  |  |
| Brodie III, 1993<br>(Experiment 1) | Coral snakes<br>(*Micrurus nigrocintus*) | Aposematism | 408 | 32 | 0.08 | Not<br>reported | - | 48 | No | Yes |
| Brodie III, 1993<br>(Experiment 2) | Coral snakes<br>(*Micrurus* spp. and others) | Mimicry | 2100 | 120 | 0.06 | 76 | 0.04 | 72 | No | Yes |
| Pfennig *et al.*, 2001<br>(Experiment 1) | Coral snakes<br>(*Micrurus fulvius*)<br>and kingsnakes<br>(*Lampropeltis triangulum*) | Batesian mimicry | 480 | 25* | 0.05 | Not<br>reported | - | 672 (4 weeks) | No | Yes |
| Pfennig *et al.*, 2001<br>(Experiment 2) | Coral snakes<br>(*Micruroides euryxanthus*)<br>and kingsnakes<br>(*Lampropeltis pyromelana*) | Batesian mimicry | 720 | 49* | 0.06 | Not<br>reported | - | 336 (2 weeks) | No | Yes |
| Vignieri *et al.*, 2010 | Beach mice<br>(*Peromyscus polionotus*) | Crypsis | 896 | 8 | 0.01 | 20 | 0.02 | 72 | No | Yes† |
| Valkonen *et al.*, 2011<br>(Experiment 1) | Viper snakes<br>(*Vipera* spp.) | Aposematism vs.<br>disruptive coloration | 900 | Not<br>reported | 0.09 | Not<br>reported | 0.07 | 44-47 | No | Yes |

**Table 3.** (Continuation)
| Publication | Organism | Hypothesis | Total<br>models | Bird marks |  | Non-bird marks |  | Exposure time<br>(h) | Colour<br>calibration | Support for<br>hypothesis |
| --- | --- | --- | --- | --- | --- | --- | --- | --- | --- | --- |
|  |  |  |  | No. | Freq. | No. | Freq. |  |  |  |
| Valkonen *et al.*, 2011<br>(Experiment 2) | Viper snakes<br>(*Vipera* spp.) | Aposematism vs.<br>disruptive coloration | 360 | Not<br>reported | 0.10 | Not<br>reported | 0.04 | 42-49 | No | Yes |
| Comeault & Noonan, 2011 | Poison frogs<br>(*Dendrobates tinctorius*) | Frequency-dependent<br>selection for<br>aposematism | 1891 | 49 | 0.03 | Not<br>reported | - | 72 | No | Yes |
| Merrill *et al.*, 2012 | *Heliconius* butterflies<br>(*Heliconius cydno* and<br>*H. melpomene*) | Frequency-dependent<br>selection for mimicry | 1440 | 58 | 0.04 | Not<br>reported | - | 72 | Yes | Yes |
| This study | *Heliconius* butterflies<br>(*Heliconius erato*) | Frequency-dependent<br>selection for mimicry | 2160 | 56 | 0.03 | 667 | 0.31 | 72 | Yes | No |

\* Predation marks were not bird marks but carnivore marks (e.g. black bears, bobcats, coyotes, foxes or raccoons)
† Hypothesis test was not only based on bird marks but also on mammals marks
NOTE: all these studies reported the occurrence of bird and non-bird marks; not all of them, however, reported non-bird marks numbers

Nevertheless, our results could have other implications that have rarely been explored. It is possible that hybrid forms may only be at a disadvantage when they differ substantially in appearance from locally occurring abundant pure forms. It is well known that predators, including birds, can generalise their response in avoiding familiar aposematic prey towards novel or unfamiliar prey of similar appearance (e.g. Aronsson & Gamberale-Stille, 2008; Ihalainen *et al*., 2012). However, how and when generalisation works is poorly understood, and research aiming to understand generalisation behaviour has generally been held back because few studies have analysed warning coloration from the perspective of the predator’s visual and cognitive systems rather than a human perspective (Stevens & Ruxton, 2012). In addition, most work has focussed on one aspect of prey appearance at a time (e.g. colour) and rarely considered the entire appearance of the prey item, which may be crucial in generalisation. It is also possible that predators use key diagnostic features in generalising between aposematic species (e.g. the presence of red and black patches along with characteristic wing shapes). Therefore, in this study the avian predators could have been very effective in generalising the appearance of the different morphs and associating key features with unpalatability. This is an area that needs much more research and could have significant implications for the evolution of mimicry and the fitness of novel colour forms as a whole. Although we did not detect a colour pattern effect on bird marks, there was a strong effect of ‘zone’, with higher attack rates observed within the hybrid zone (Figure 4B). This observation offers a possible insight into the behaviour of bird predators when exposed to a diversity of colour patterns. It has been shown experimentally that the effectiveness of aposematic signals decreases when a larger diversity of signals is available to a bird predator (Ihalainen *et al*., 2012). In the hybrid zone, there are at least 10 hybrid patterns of *H. erato* and at over 10 more of *H. melpomene*. This large diversity of aposematic signals can potentially explain the lower survival of all tested colour patterns in the hybrid zone, because the predator bird community may have a harder time learning all these different patterns and identifying them all as distasteful, and attack them indiscriminately.

Finally, an additional important effect detected was a random effect of the variation between experimental sites on non-bird marks (Figure 5B, Table2). This effect was explained by anecdotal observations about the local abundance of some non-bird ‘predators’, notably sweat bees and grasshoppers, which varied widely among experimental sites. Thus, it is clear that accidental (non-visual) predators, such as insects, attack the butterfly models randomly, without respect to colour pattern or zone.

### Do birds target the head?

Our data clearly support the hypothesis that birds attack the head section of the body, rather than any section at random. Other studies have also found a tendency for birds to attack head-like structures. For instance, in a large sample of wild-caught Lycaenids, Robbins (1981) found that species with a conspicuous false-head, in the anal angle of the hind-wings, showed five times more damage from predator attacks on their hind-wings than species that lack the false-head. Likewise, Wourms & Wasserman (1985) showed experimentally that head-like structures, painted on the hind-wings of a butterfly that naturally lacks a false-head, increased the chances that blue jays would attack the hind-wing area, away from the real head. Furthermore, Wourms & Wasserman (1985) also found that the deflected attacks increased the chance that the butterflies would escape during the necessary prey handling that happens after the attack. In both studies, the head-like structures that fooled predators included features that are suggestive of a butterfly head (e.g. false antennae or eye-spots). In our study, however, the plasticine bodies did not have any feature that would suggest that one of the ends was meant to be the head (Figure 1). Therefore, in our models birds must recognize the head section by the orientation, as suggested by wing shape and colour pattern. It can be argued that the metallic reflection of the pin, placed on the boundary between the anterior and middle sections, could have attracted birds to this area. Even though we cannot rule out this possibility, the fact that the frequency of bird marks that affected the most anterior section only, without affecting the pin area, is almost as large as the frequency of marks that affected the pin (23.2% vs. 37.5%), suggests that the pin did not influence the body section targeted by birds. These results also strongly indicate that our artificial prey were effective: birds attacked our models as if they were real butterflies.

### On the use of artificial *Heliconius* to study bird predation and natural selection

In spite of our negative results with respect to colour pattern effects, we still recommend the use of artificial *Heliconius* to study bird predation on these butterflies. In fact, a recently published study has succesfully used *H. erato* models to detect lower predation rates over butterflies that roost gregariously compared to butterflies that roost solitarily (Finkbeiner *et al*. 2012). Furthermore, another study, performed in Panama around the same time as the study here reported, succeeded in detecting differences in bird predation associated with colour pattern, using models constructed in exactly the same way as those used in our experiment (Merrill *et al*. 2012).

In general, studies with artificial prey are complementary to mark recapture experiments, such as those performed by Benson (1972), Mallet & Barton (1989) and Kapan (2001), because artificial prey provide different technical advantages. For instance, predators can be easily recognized by the marks that they leave in the models, which allow researchers to keep a record of predation events, instead of just inferring these events from individual disappearances. Also, models allow researchers to test for the effects of morphological characters, such as colour pattern or body shape, while controlling for other potentially confounding factors. In the present study, for instance, it was necessary to control for the possible effects of adaptations associated with the altitudinal gradient across the hybrid zone (e.g. variation in thermal tolerance). Finally, there is usually little constraint in the number of models that can be exposed to predation, and therefore it is possible to attain larger sample sizes than those attained when using live specimens. Despite these advantages, there are a few considerations that we suggest should be taken into account in future studies. In particular, *Heliconius* aposematic patterns are likely to reduce the chances of the models being attacked, increasing the effort required to gather a large enough sample of bird marks. It seems important, in retrospect, to focus on gathering a large number of predator marks rather than favouring the total number of models exposed to predation. In the present study, for instance, replacing models that were accidentally marked by non-bird ‘predators’—such as ants, grasshoppers or sweat bees—rather than taking these models away from the experiment, would have probably increased the chances of recording more bird marks. A further desirable addition would be to include non-aposematic (either palatable or fake) colour-patterns as control treatments, since they provide information about overall rates of attack and the degree of aposematism of the models, and/or whether the artificial prey appear realistic to predators.

Overall, our study has shown that the use of artificial *Heliconius* is helpful in exploring aspects of bird predatory behaviour for which there is little evidence of their importance in the field. In particular, the effects of prey community structure on birds’ discriminatory behaviour that selects for Müllerian mimicry had been elegantly demonstrated in captivity (Ihalainen *et al*., 2012), but not backed up by field observations yet. Our discovery that the rate of bird attacks is substantially larger in a morphologically diverse prey community, such as a hybrid zone, is an important piece of evidence in support of the idea that a complex aposematic prey community can impact the patterns of bird predation in nature, and—as a consequence—can impact the evolution of Müllerian mimicry as well. According to Ihalainen *et al*.’s (2012) experiment in the lab, accurate mimicry is more likely to evolve in simpler prey communities. Hence, our study has opened new opportunities to address questions about the interactions between *Heliconius* butterflies and their bird predators.

## ACKNOWLEDGEMENTS

We would like to thank Richard Merrill and Steven Shaak for their collaboration during the colour calibration of *Heliconius* models. Nicola Nadeau and Agata Surma assisted in the field during the pilot experiments, while Carlos Robalino and Elizabeth Heras did so during the large-scale experiment. Jens Töniges and Robby Flemisch, from the Curiquingue Foundation (Tena), and Carmen Luzuriaga, from the Pindo-Mirador Biological Station (Universidad Tecnológica Equinoccial), provided great logistical support while we worked in their field stations. Santiago Villamarín and Marco Altamirano, from the Ecuadorian Museum of Natural Sciences, sponsored and helped us in obtaining research, collection and export permits from the Ecuadorian Ministry of Environment. The Ministry of Environment, in turn, through its provincial offices in Napo and Pastaza, provided us with the aforementioned permits. Finally, Patricia Salazar provided constant logistical support during the whole fieldwork period in Ecuador. Financial support came primarily from a Leverhulme Trust award to C.D.J., the University of Cambridge Trusts to P.A.S, the Balfour Fund (Zoology Department at Cambridge) to I.O. MS was supported by a Biotechnology and Biological Sciences Research Council David Phillips Research Fellowship (BB/G022887/1).

